# The *Plasmodiophora brassicae* effector PbEGF1 manipulates plant immunity and regulate primary infection

**DOI:** 10.1101/2024.07.23.604797

**Authors:** Hui Yang, Yushu Xu, Yushan Zhao, Yinping Shu, Xin Sun, Junbo Du

## Abstract

*Plasmodiophora brassicae* causes a significant global threat to cruciferous vegetables and crops. However, the current comprehensions of its pathogenic ways is still unclear. This study identified a *P. brassicae* effector, called PbEGF1, which strongly induces cell death in *N. benthamiana.* Notably, *PbEGF1* was significantly up-regulated in seedlings inoculated with highly virulent *P. brassicae*, indicating a pivotal role for PbEGF1 in pathogenicity. Furthermore, overexpression of PbEGF1 in hosts enhanced susceptibility to *P. brassicae,* and promoted elongation of root hairs, thus creating favorable conditions for root hair infection. Silencing of *PbEGF1* reduced the pathogenicity of *P. brassicae*. This finding confirms the significance of primary infection in host recognition and interaction with *P. brassicae*. To further elucidate the virulence function of PbEGF1, we identified BnNHL13 (nonrace-specific disease resistance 1/harpin-induced 1-like 13) as its target protein. Silencing *BnNHL13* enhanced host susceptibility to *P. brassicae,* and promoted root hairs elongation, indicating that down-regulation of *BnNHL13* was more conducive to establishing *P. brassicae* infection. Subsequent investigation revealed that PbEGF1 has the ability to induce degradation of the BnNHL13 protein, thereby disrupting the host defense response and facilitating *P. brassicae* infection. Our findings provide novel insights into genetic strategies for enhancing plant resistance against clubroot disease.

## Introduction

The devastating clubroot disease, caused by the obligate biotrophic parasite *Plasmodiophora brassicae*, has a global impact on cruciferous vegetables and crops. Up to now, controlling clubroot disease remains challenging, emphasizing the necessity for an improved understanding of *P. brassicae* biology.

The completion of the *P. brassicae* life cycle relies on a susceptible host (Kageyama and Asano, 2009). Within the soil, resting spores germinate and subsequently release primary zoospores to infect root hairs and epidermal cells, constituting a stage commonly referred to as primary infection or root hair infection. Subsequently, *P. brassicae* forms zoosporangia that release secondary zoospores, initiating a secondary infection in the root cortex, which ultimately leads to the development of long-lived resting spores. Several studies have discovered the role of root hair infection (primary infection) and cortical infection (second infection). It has always been believed that the secondary infection of *P. brassicae* plays a significant role. However, several subsequent studies have gradually demonstrated the significant role of primary infection. For instance, inoculation solely with secondary zoospores led to a diminished disease severity in susceptible hosts compared to primary zoospores infection (McDonald *et al*., 2014). This finding suggest that the host’s recognition of infection and subsequent initiation of a host response predominantly occur during the primary infection. To date, it remains unclear how *P. brassicae*’s primary infection interferes with host growth or immunity to facilitate subsequent infection.

Plant pathogens have evolved a diverse range of effectors, which can deliver into host cells, either suppressing plant immunity or inducing changes in plant growth (Presti *et al*., 2015). Similar to other plant pathogens, *P. brassicae* possesses secreted proteins and exhibits effector-like characteristics. The secretory activity of certain putative effectors in *P. brassicae* has been experimentally validated (Schwelm *et al*., 2015; Rolfe *et al*., 2016; Chen *et al*., 2019; Pérez-López *et al*., 2020). Recent studies have also identified specific effectors during the early and late stage of infection, as well as differentially expressed effectors associated with primary and secondary infection (Chen *et al*., 2019; Pérez-López *et al*., 2020; Yang *et al*., 2022). Effector secretion serves as a strategy employed by pathogens to interfere with host growth processes. Studying gene function and pathogenesis in *P. brassicae* presents challenges compared to fungi, bacteria, and oomycetes due to the unavailability of a transformation system. Up to now, a limited number of effectors function have been reported for *P. brassicae*. It is still unclear how *P. brassicae* induces aberrant root growth to facilitate its own infection.

Compared to other agriculturally significant pathosystems, the interactions mechanism between Plasmodiophora and its host remains limited. This study identified a *P. brassicae* effector, called PbEGF1, which strongly induces cell death in both host and non-host plants. The overexpression of PbEGF1 facilitates the colonization of *P. brassicae* in host root hairs and enhances their growth. Furthermore, we identified a protein designated as BnNHL13 (nonrace-specific disease resistance1/ harpin-induced-like13, NDR1/HIN1-like 13) from *B. napus*. NHL13 has been reported to be involved in the activation of innate immunity and plant growth in *Arabidopsis thaliana* (Xin *et al*., 2015). BnNHL13 serves as the target protein for the *P. brassiace* effector PbEGF1. In this study, our results highlight a counter strategy employed by *P. brassicae* to promote infection by disrupting NDR1-mediated responses. Revealing the pathogenic strategy is critical for breeding clubroot-resistant plant variety.

## Materials and methods

### Resting spores preparation and inoculation

XC isolates of *P. brassicae* were obtained from Xichang in Sichuan Province, China, and were identified as pathotype 4. SF isolates was obtained from Shifang in Sichuan Province and were classified as pathotype 11 using the differential system of Williams. CD isolates of *P. brassicae* were obtained from Chengdu in Sichuan Province, China, and were identified as pathotype 4. Resting spores of *P. brassicae* were prepared following the method described (Ji *et al*., 2014). The concentration of resting spores was adjusted to 1×10^7^ spores/ml for inoculation, a volume of 300 µL of the spore suspension was applied around the root zone of each plant.

### Morphological characterization of P. brassicae

After inoculating the seedlings with *P. brassicae* for approximately 7-14 days, the plants were uprooted to remove any soil adherence from the roots. Subsequently, the roots were stained with 0.5% fluorescent pink dye for approximately 1-2 minutes, subsequently removing any background coloration through sterilized water treatment. This staining facilitated the observation of *P. brassicae* zoosporangia, plasmodia and resting spores within the root hair and epidermal cells under microscope (Carl Zeiss, Germany).

### Secretory activities assessment for PbEGF1 signal peptides in yeast

The high-fidelity DNA polymerase was used to amplify the signal peptide of *P. brassicae* effector PbEGF1. The primer sequences are provided in Supplementary Table 1. Subsequently, the purified DNA fragments were individually ligated into the pSUC2 vector and transformed into the yeast strain YTK12. Secretory activity was assessed through growth assays following the protocol described (Jacobs *et al*., 1997). The YTK12 strains exhibiting secretory activity were capable of growth on YPRAA medium (2% peptone, 1% yeast extract, 2 µg/mL antimycin A and 2% raffinose) and CMD-W medium (2% sucrose, 0.1% glucose, 0.67% yeast N base without amino acids, 0.075% tryptophan dropout supplement and 2% agar). Invertase activity was assessed based on the enzymatic conversion of colorless 2, 3, 5-triphenyltetrazolium chloride (TTC) to insoluble red-colored 1, 3, 5-triphenyl-formazan (TPF).

### PbEGF1 transient expression in Nicotiana benthamiana, Arabidopsis thaliana and Brassica rapa leaves

High-fidelity DNA polymerase was used to amplify the full-length nucleotide sequence of PbEGF1. Primers were designed based on the sequence of *P. brassicae* PBRA_ 007830, accession number CDSF01000097.1. Subsequently, the PCR products were ligated into the pBIB-BASTA-35S-GWR-GFP (Green fluorescent protein) vector. The primer sequences are provided in Supplementary Table 1. The recombinant vector was then transformed into *Agrobacterium tumefaciens* strain GV3101, which then cultured until centrifugation. The *A. tumefaciens* cells were resuspended in MMA infiltration solution containing 10 mM MgCl_2_, 10 mM MES and 150 mM acetosyringone. At the four-leaf to six-leaf stage, the prepared solution (optical density at 600 nm [OD600]=0.5) was infiltrated into leaves of *N. benthamiana, A. thaliana* (Col ecotype) and *B. rapa* using a needleless syringe to overexpress PbEGF1.

### Oxygen burst detection

*A. tumefaciens* cultures were infiltrated into the leaves of *N. benthamiana*. Subsequently, at one post-infiltration, the leaves were excised. The detection of reactive oxygen species in *N. benthamiana* leaves was conducted using 3,3’-diaminobenzidine stain, following a previously described protocol (Hans *et al*., 1997).

### Host induce gene silencing and transient overexpression of PbEGF1 in host roots

The 312-bp partial forward fragment of PbEGF1 was inserted into the XhoI/KpnI site of the pKANNIBAL vector, followed by the insertion of the corresponding reverse fragment into the HindⅢ/Xba Ⅰ site of the same vector. The primer sequences are provided in Supplement Table 1. Both fragments in the pKANNIBAL vector contain a BsrGI restriction enzyme recognition site. Subsequently, these fragments were cleaved by BsrGI and inserted into the corresponding site of the pBIB-BASTA-35S-GWR-GFP vector, named PbEGF1-HIGS. Finally, the recombinant vectors were transferred into the *A. tumefaciens* strain GV3101, and the transformed strains were cultured, centrifuged, and resuspended in MMA infiltration solution.

We employed susceptible *Brassica napus* cultivars Chuanyou 36. The aforementioned MMA solution was used to immerse seeds, which were gently agitated for a duration of 15-20 hours (Zhang *et al*., 2023; Yang *et al*., 2024). Subsequently, the seeds were sown in soil under controlled conditions at a temperature of 25℃ with a light/dark cycle of 16 hours and 8 hours, respectively. A volume of 500 μl of the aforementioned MMA solution was applied to the soil surrounding the roots of seven-day-old seedlings. The roots from twenty-one-day-old seedlings were collected to assess the efficiency of silencing.

The ten-day-old seedlings were used for the inoculation of *P. brassicae,* a 300 μl suspension of resting spores was carefully applied to the soil surrounding the root of each seedling. Disease severity index (DSI) was subsequently assessed at 25-30 dpi using a standardized scale ranging from 0 to 3 (Siemens *et al*., 2002). Statistical analysis was performed using a minimum of 50 seedlings.

### Fluorescence observation was conducted in the host roots and P. brassicae

The seeds were gently agitated for 24 hours with MMA solution containing transformed *A. tumefaciens* GV3101 carrying 35S-PbEGF1-GFP and 35S-PbEGF1-HIGS-GFP, respectively. Subsequently, the seeds were placed in petri dishes. Three-day-old seedlings were used for fluorescence analysis. Ten-day-old seedlings were inoculated with *P. brassicae.* Then, the seedlings were treated with the aforementioned MMA solution at 4 dpi. At 7 dpi, the roots of seedlings were taken to observe fluorescence under a Carl Zeiss microscope.

### Quantitative real-time reverse transcription-polymerase chain reaction

*Brassica napus* variety Chuanyou 36 is susceptible to the CD, XC and SF isolates of *P. brassicae*. *B. rapa* variety T1-145 is susceptible to XC isolates but tolerant to SF isolates. Root samples of *B. rapa* T1-145 were collected at 4, 10, 16, and 45 dpi. Total RNA extraction from *N. benthamiana* leaves, *B. napus,* and *B. rapa* roots was performed according to previous method (Yang *et al*., 2024). For quantifying the relative expression levels, the housekeeping gene actin was used as an internal control in *B. rapa*, *B. napus*, *N. benthamiana*, or *P. brassicae*, respectively. The 2^-ΔΔCt^ method was employed to determine the relative expression levels, with three replicates performed for each analysis (Livak *et al*., 2001). The primer sequences was provided in Supplementary Table 1.

### Yeast two-hybrid interaction assays

The cDNA sequences of PbEGF1 full-length were cloned and ligated into the PGBKT7 vector. Strains carrying the PGBKT7-PbEGF1 vector were co-cultured with the *B. napus* root library strain, and the target protein of PbEGF1 in the host was identified using a yeast two-hybrid (Y2H) analysis. Subsequently, we focused on investigating the interaction of BnNHL13 as a potential target protein. The cDNA sequences of BnNHL13 full-length were cloned and ligated into the PGADT7 vector. Primer sequences are listed in Supplement Table 1. The PGADT7-BnNHL13 and PGBKT7-PbEGF1 recombinant vectors were co-transformed into Y2H gold competent yeast cells. These cells were cultivated in SD/-LEU/-TRP (DDO) medium for 72-96 hours at 30℃. The correct monoclonal colonies were selected, they were resuspended in sterile water and then cultured on SD/-ADE/-HIS/-LEU/-TRP/ X-a-Gal/AbA (QDO/A/X) medium at 30℃ for 72-96 hours.

### Co-immunoprecipitation analyses

*N. benthamiana* leaves were infiltrated with *A.tumefaciens* GV3101 to achieve a final optical density of OD600 = 0.5. Samples were collected at 36 hours post-infiltration. Proteins from these samples were extracted using lysis buffer and subsequently subjected to centrifugation at low temperature (4℃) and high speed (13,000×g) for fifteen minutes. The supernatants obtained from the samples were incubated with 10 µL anti-GFP magnetic beads (Chromotek, Germany) for three hours under gentle agitation at 4℃. The beads were subsequently pelleted through centrifugation at 2,500×g and then washed five times. The protein complexes were eluted, and subsequent denaturation was employed using a loading buffer containing sodium dodecyl sulfate (SDS) at a five-fold concentration to boil for fifteen minutes. Protein analysis was performed using western blotting.

### Validation of luciferase complementation

The PbEGF1 construct was generated by fusing it to the 35S-LUS-N, and the predicted target protein BnNHL13 was fused to the 35S-LUS-C. The primer sequences are listed in Supplement Table 1. Subsequently, both recombinant vectors were transformed into *Agrobacterium* GV3101. The GV3101 solution was then infiltrated into *N. benthamiana* leaves and cultured under standard greenhouse conditions for 24-48 hours. After the infiltration, the backside surface of the leaves was uniformly moistened with a reaction solution containing a 1 mM concentration of D-luciferin substrate. Afterwards, *N. benthamiana* were placed in darkness for 10 minutes before their leaves were excised and positioned upside down in petri dishes. Finally, these leaf samples were transferred to a plant imaging system for luminescence measurement.

## Results

### PbEGF1 triggers cell death in N. benthamiana and host plants, as well as induces a plant immunity response

In a previous study, candidate effectors were identified based on the prediction of potential secreted proteins with an N-terminal signal peptide and the absence of transmembrane domain. These selected effectors with signal peptides were subsequently cloned into a plant expression vector, which added a C-terminal GFP tag to the proteins. Transient expression in *Nicotiana benthamiana* has been commonly employed to investigate the effects of pathogen effectors such as *Phytophthora sojae* and *Bursaphelenchus xylophilus* on plant defense (Ma *et al*., 2015; Hu *et al*., 2019). These vectors were then transformed into *Agrobacterium tumefaciens* GV3101 for transient expression in *N. benthamiana*. Among them, two genes exhibited significant induction of cell death. Besides Pb257 (Yang *et al*., 2024), another effector containin an EGF domain was identified using the Simple Modular Architecture Research Tool (SMART) and designated as PbEGF1 for further investigation.

Transient expression of PbEGF1 in *N. benthamiana* leaves strongly induced cell death at five days post-infiltration (Fig. 1A). Additionally, PbEGF1 transient expression induced a significant elevation in reactive oxygen species levels within the infiltrated leaves at one day post-infiltration (Fig. 1B), as well as up-regulation expression of defense-related genes including *NbMKK4*, *NbMPK3*, *NbPR1*, and *NbNPR3* (Fig. 1C), thereby demonstrating its influence on plant defense pathways.

**Fig. 1.**
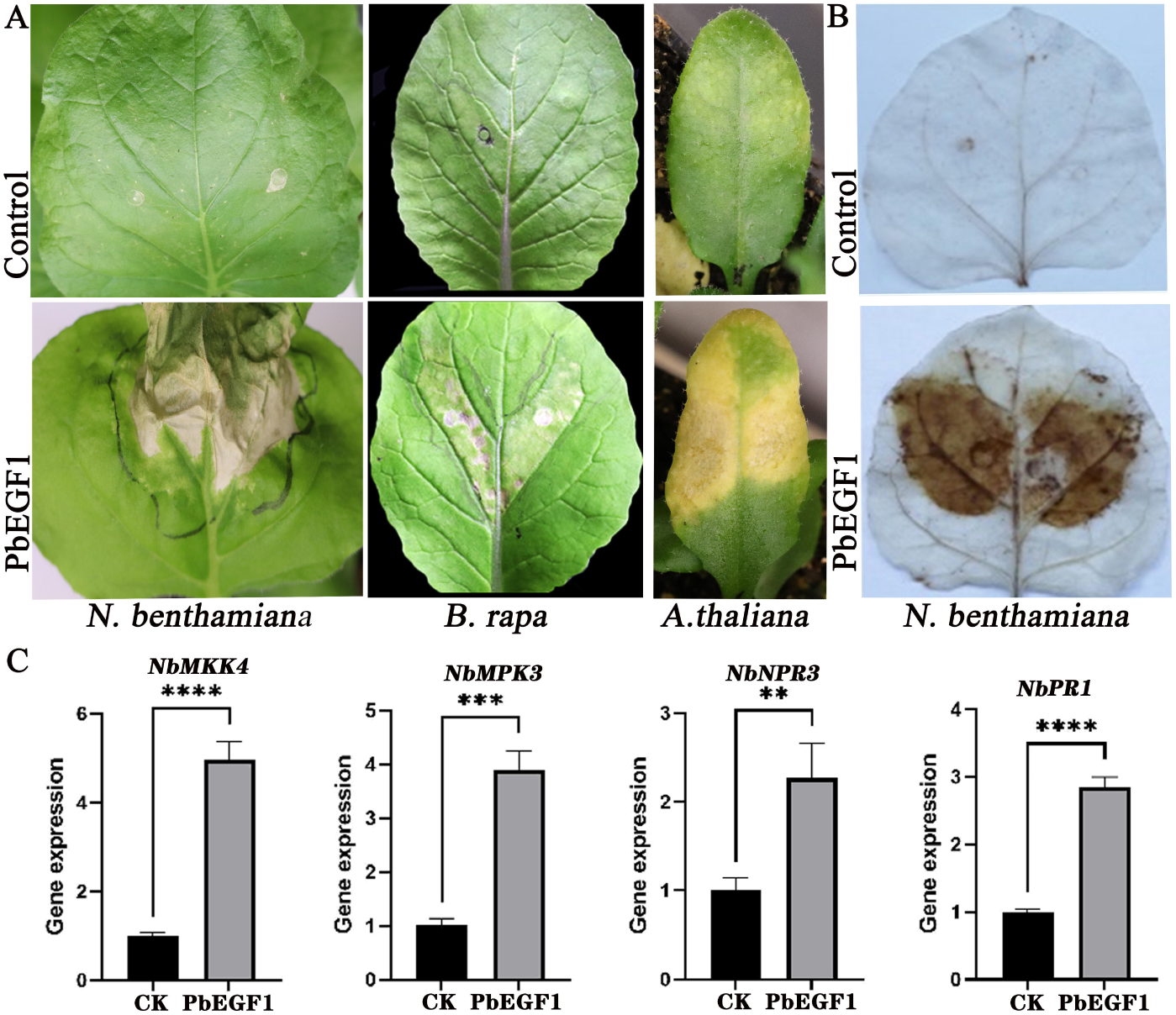
The putative effector PbEGF1 of *P. brassicae* induces cell death and a burst of reactive oxygen species (ROS). A, Infiltration of PbEGF1 into leaves induces cell death in *N. benthamiana*, host *A. thaliana* ecotype Col and *B. rapa* at 5 days post-infiltration. B, Infiltration of PbEGF1 into leaves triggers a rapid burst of reactive oxygen species (ROS) in *N. benthamiana* at one day post-infiltration. C, Real-time fluorescent quantitative PCR (qPCR) analysis reveals the up-regulation expression of defense genes in *N. benthamiana* leaves at 3 days post-infiltration with PbEGF1, compared to the negative control GFP infiltration (Control). The experiment was repeated three times for statistical significance assessment using one-way ANOVA followed by Tukey’s multiple comparison test (**p<0.01, ***P<0.001).

Furthermore, transient expression of PbEGF1 also induced cell death in host *Arabidopsis thaliana* Col ecotype and *B. rapa* leaves (Fig. 1A). These findings indicate the ability of multiple plant species to recognize and respond to PbEGF1.

### PbEGF1 triggers cell death in N. benthamiana when secreted into the apoplast

The open reading frame of PbEGF1 encodes a 238-amino acid protein. PbEGF1 contains a signal peptide (SP) consisting of the first 19 amino acids at its N-terminus (Fig. 2). To confirm its secretory function, a SP trap system was utilized in yeast. Yeast strains expressing SUC2 fused with the PbEGF1 signal peptide enabled growth of YTK12 on both CMD-W and YPRAA medium (Fig. 2). Moreover, transformed yeast strains exhibited invertase activity, which efficiently converted colorless TTC into insoluble red TPF (Fig. 2). These results indicate that the signal peptide of PbEGF1 is functional in yeast, and confirm that secretion of PbEGF1 occurs via the classical pathway.

**Fig. 2.**
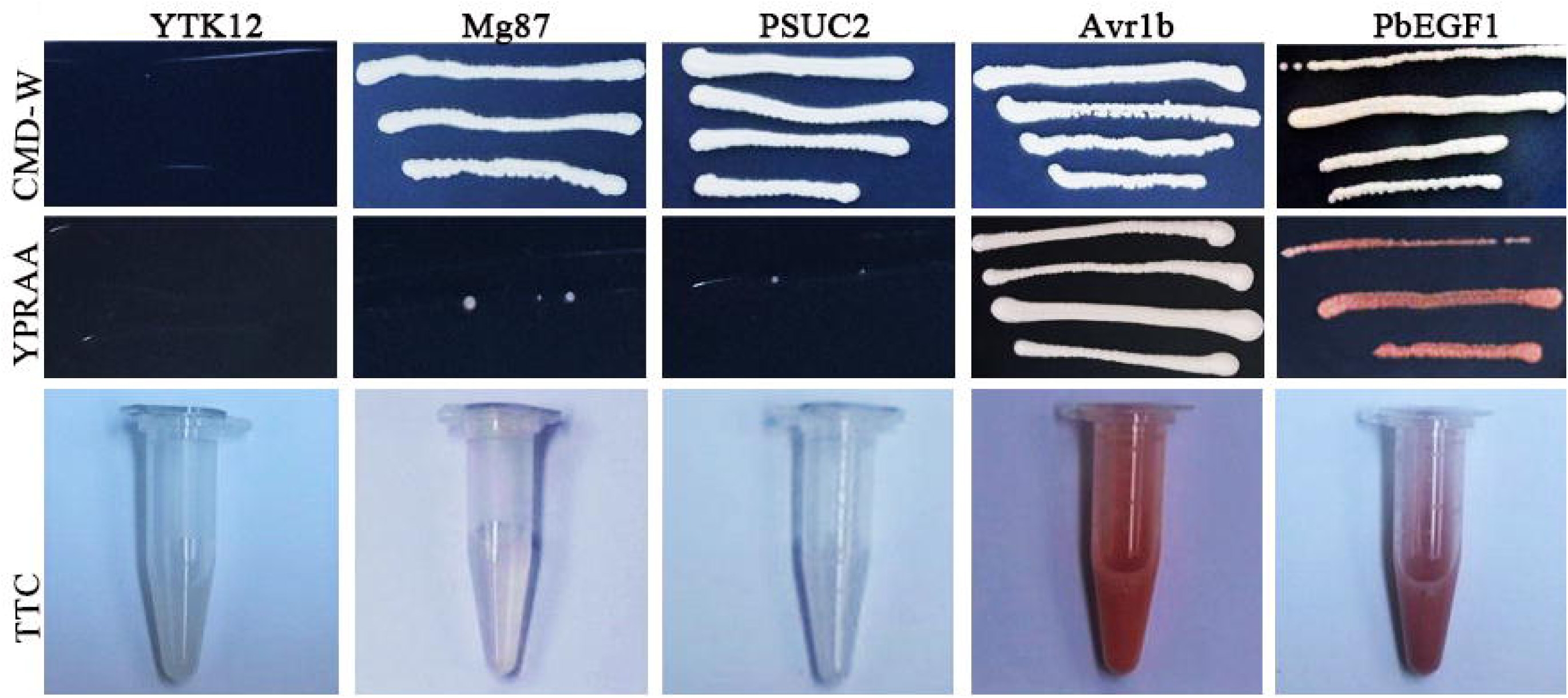
The function of the PbEGF1 signal peptide was validated using a yeast invertase secretion assay. The predicted sequence of PbEGF1 signal peptide was fused in-frame with the yeast invertase sequence in the pSUC2 vector. This functional signal peptide enabled transformed YTK12 yeast to grow on YPRAA medium supplemented with raffinose as the sole carbon source. The viability of transformed yeast strains grown on CMD-W medium was consistent across three replicates of this experiment. Untransformed YTK12 yeast did not exhibit any growth on either CMD-W or YPRAA media. Positive control sequences from P. sojae Avr1b and negative control sequences from M. oryzae Mg87 were employed.

To investigate the role of the PbEGF1 signaling peptide in triggering cell death, we transiently expressed PbEGF1 lacking the signal peptide (PbEGF1^nsp^) in *N. benthamiana* leaves. However, PbEGF1^nsp^ failed to elicit cell death. Otherwise, it has been proven that pathogenesis-related (PR1) protein is predominantly localized in the apoplast (Van Loon, 1985). To determine whether PbEGF1 was secreted into the apoplast to trigger cell death, we deleted its N-terminal signal peptide and fused it with the signal peptide from PR protein 1 (PR1) of *N. tabacum* to produce PR1^SP^-PbEGF1 (Fig. 3A). In transient assays, both PbEGF1 and PR1^SP^-PbEGF1 were found to trigger cell death in *N. benthamiana* (Fig. 3A). Western blot analysis confirmed the expression of all proteins in *N. benthamiana* leaves (Fig. 3C). These results indicate that PbEGF1 triggered cell death in *N. benthamiana* when it secreted into the apoplast.

**Fig. 3.**
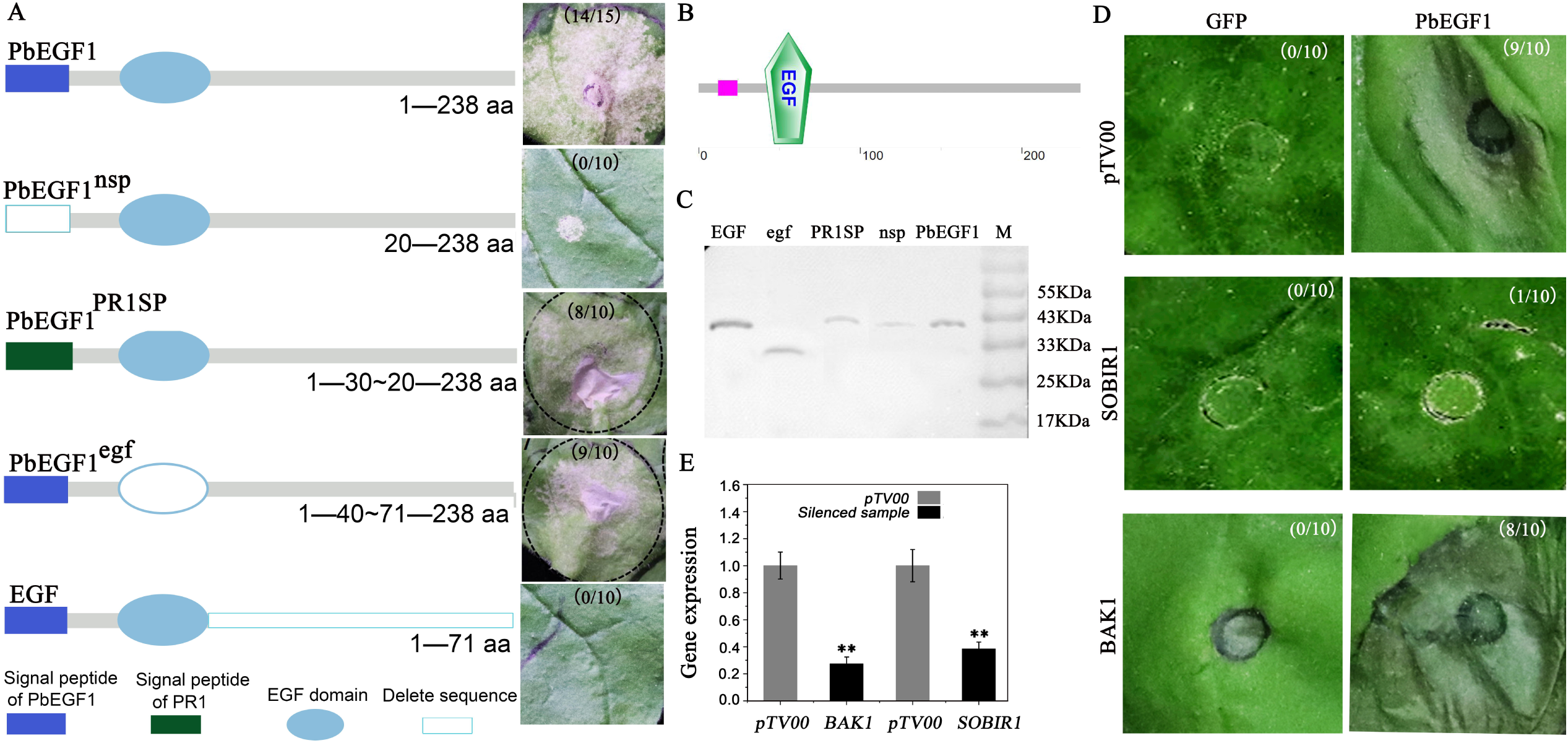
The presence of PbEGF1 signal peptide as well as NbSOBIR1 in *N. benthamiana* is essential to trigger cell death. A, a series of truncation mutants were generated and transiently expressed via agroinfiltration in *N. benthamiana* leaves to investigate their effects on inducing cell death. Photographs were taken at five days after infiltration. B, Protein sequence analysis of PbEGF1 was conducted using the Simple Modular Architecture Research Tool (SMART). C, Western blotting was performed to detect the expression of PbEGF1 and its truncation mutants in infiltrated leaves. EGF refers to a sequence containing both a signal peptide and an EGF domain, egf refers to a sequence without an EGF domain. PbEGF1 represents the complete sequence, nsp represents the PbEGF1 sequence without the signal peptide, and PbEGF1^PR1SP^ refers to the fusion between the signal peptide of PR protein 1 and the PbEGF1 sequence. D, Representative images of PbEGF1-induced cell death in *N. benthamiana* leaves with silenced pTV00 (control), *BAK1*, or *SOBIR1*. GFP was used as a control proteins. *Agrobacterium* carrying PbEGF1 was infiltrated into the leaves 14 days after infiltration of *Agrobacterium* carrying *BAK1* or *SOBIR1* silencing vector. E, qRT-PCR analysis was performed to determine the transcript levels of *SOBIR1* and *BAK1*. The experiment was repeated three times for statistical significance assessment using one-way ANOVA followed by Tukey’s multiple comparison test (**p<0.01).

PbEGF1 possesses a typical EGF domain. To analyze the role of this domain in inducing cell death, PbEGF1 lacking EGF domain for amino acids 41-70 (PbEGF1^egf^), as well as the EGF domain from amino acids 1-71 (EGF) was separately constructed into the plant vectors. *Agrobacterium*-mediated expression of PbEGF1^egf^ successfully induced cell death in *N. benthamiana* leaves. Relevantly, over-expression of the EGF domain (EGF) failed to elicit cell death (Fig. 3A). The result indicates the presence of the EGF domain is not essential for inducing cell death.

### NbSOBIR1 but not NbBAK1, is required for PbEGF1-induced cell death in N. benthamiana

BAK1 and SOBIR1 can recognize pathogen-associated molecular patterns (PAMPs). Previous studies demonstrated the ability of PAMP INF1, secreted by *Phytophthora infestans*, and RcCDI1, derived from *Rhynchosporium commune*, to induce cell death in the apoplast of *N. benthamiana* through the involvement of NbBAK1 and NbSOBIR1 receptors (Franco-Orozco *et al*., 2017; Heese *et al*., 2007). It remains unclear whether PbEGF1 functions as a PAMP and if NbBAK1 or NbSOBIR1 is involved in PbEGF1-induced cell death. Therefore, we employed virus-induced gene silencing (VIGS) to silence *NbBAK1* or *NbSOBIR1* individually in *N. benthamiana*. The silencing efficiency of both *NbBAK1* and *NbSOBIR1* was validated by qRT-PCR analysis. The silencing plants showed decreased expression for either *NbBAK1* or *NbSOBIR1* compared to pTV00 control plants (Fig. 3E). We employed VIGS to individually silence *NbBAK1* or *NbSOBIR1* while simultaneously expressing *PbEGF1* in *N. benthamiana*. Silencing of *NbSOBIR1* failed to induce cell death upon expression of *PbEGF1* (Fig. 3D). However, leaves with silenced *NbBAK1* exhibited cell death upon transient expression of *PbEGF1* (Fig. 3D). These results indicated that NbSOBIR1 is involved in mediating PbEGF1-triggered cell death.

### Diverse expression patterns of PbEGF1 during the infection stage of P. brassicae strains with different levels of virulence

In view of the impact of PbEGF1 on plant defense, we further investigated the expression pattern of PbEGF1 during *P. brassicae* infection to assess its role in pathogenicity. Previous studies have identified highly and moderately virulent strains of *P. brassicae* (Peng *et al*., 2016). To explore the potential roles of PbEGF1 in the virulence of this pathogen, we examined its expression level at different stages of *P. brassicae* infection caused by high or moderate virulence isolates (XC and SF isolates). *B. rapa* T1-145 was used as the host. The XC isolates induced typical spindle gall and exhibited high virulence, while the SF isolates mainly resulted in small spheroid galls (SSGs) and exhibited moderate virulence (Fig. 4A). Subsequently, we performed qRT-PCR analysis to evaluate the expression levels of PbEGF1. To investigate whether the expression of PbEGF1 was affected by *P. brassicae* quantity, another effector, Pb257, was selected for comparison (Yang *et al*., 2024). The internal reference for assessing effector gene expressions was *P. brassicae actin*.

**Fig. 4.**
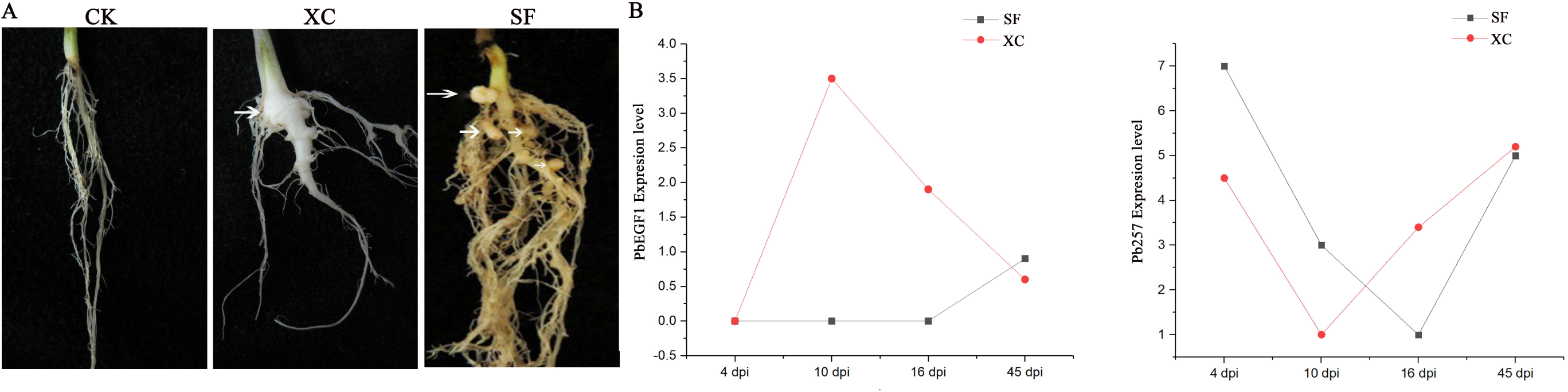
Morphology differences of T-145 plants. A, The healthy root (CK), the root infected with SF isolates resulting in small spindle galls (XC), the root infected with XC isolates leading to small spheroid galls (SF). B, *PbEGF1* expression levels were assessed via qRT-PCR in *B. rapa* T1-145 seedlings inoculated with XC or SF isolates during different infection stages.

The result revealed no significant induction of *PbEGF1* expression at 4 days post-inoculation (dpi) but reached a peak at 10 dpi in plants infected with XC isolates, subsequently, its expression level slightly decreased at 16 dpi and became low at 45 dpi. Conversely, no significant up-regulation in *PbEGF1* expression was observed in seedlings infected with SF isolates at 4, 10, and 16 dpi, however, a slight increase in expression was observedat 45 dpi, without any difference between SF and CX isolate-infected plants (Fig. 4B). These findings suggest that PbEGF1 may influence the virulence of *P. brassicae*.

It is different from *PbEGF1* expression, the expression level of *Pb257* remained consistent throughout all four time points of XC and SF infection, showing an initial increase, and a subsequent decrease, and ultimately displaying an upward trend at 45 dpi (Fig. 4B). The observation suggests that PbEGF1 expression was independent of fluctuations in *P. brassicae* quantity.

### PbEGF1 is required for full virulence of P. brassicae

The significant up-regulation of *PbEGF1* in highly virulent strains suggests its role in the pathogenicity of *P. brassicae*. However, due to its obligatory parasitism, achieving gene overexpression or knockout within *P. brassicae* genome is challenging. To overcome this limitation, we developed a method that utilizes agrobacterium-mediated transient expression and host-induced gene silencing (HIGS) techniques in *B. napus* to alter the transcript level of *PbEGF1* during *P. brassicae* infection.To design RNAi constructs capable of effectively silencing *PbEGF1*, forward and reverse DNA fragments corresponding to *PbEGF1* were individually inserted into the same vector to generate a hairpin structure with dsRNA sequence. Additionally, both the RNAi vector and overexpression vector for *PbEGF1* were separately fused with green fluorescent protein as reporter genes. The plant expression vector or RNAi vector for *PbEGF1* was transformed into agrobacterium, followed by 24-hour treatment of rapeseed seeds with agrobacterium suspension, which was prepared using MMA solution. Subsequently, the treated seeds were either sown in soil or on petri dishes.

*P. brassicae* solely infects the roots of plant. Therefore, the roots of the seedlings were excised for subsequent observation. The presence of green fluorescence in root hairs and upper epidermal cells indicated that *Agrobacterium tumefaciens* could transiently express genes in roots (Fig. 5A). Additionally, ten-day-old seedlings were inoculated with *P. brassicae* and then treated with the above agrobacterium solution at 4 dpi. The roots were observed at 7 dpi, obvious fluorescent areas were photographed (Fig. 5B), where green fluorescent indicated that primary zoosporangia can uptake expression products from its host plant.

**Fig. 5.**
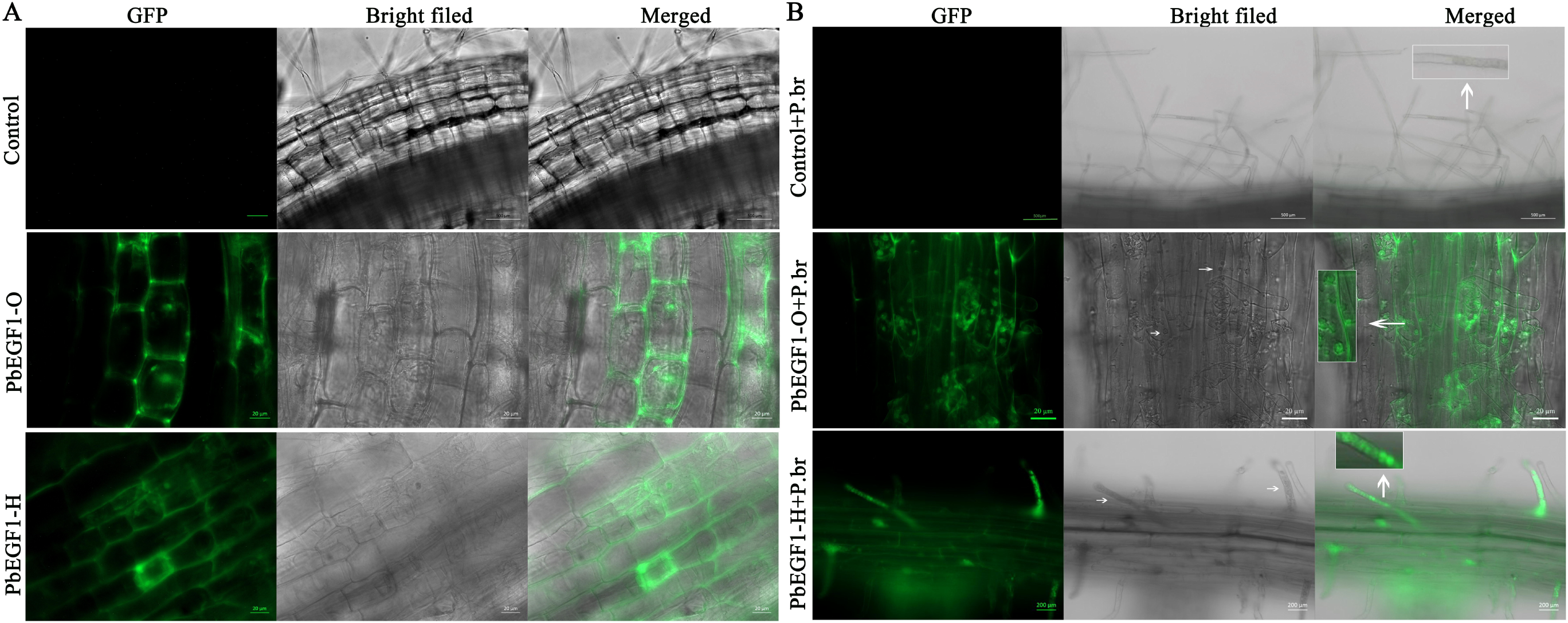
The expression of PbEGF1 in the roots of *Brassica napus* was investigated. A, Green fluorescence was observed in root hairs and epidermal cells. The PbEGF1 coding sequence was heterologously expressed and fused with green fluorescent protein (GFP) within a vector, creating a recombinant construct (PbEGF1-GFP). Additionally, a hairpin structure sequence of PbEGF1 was also fused with GFP within the vector (PbEGF1-HIGS). Three-day-old seedling were taken, fluorescence microscopy was utilized for visualization. B, Green fluorescence was observed in root cells and primary zoosporangia of *P. brassicae*. Ten-day-old seedlings were inoculated with *P. brassicae*. In addition to the seeds treated with *Agrobacterium* carrying the gene, the seedlings were treated with agrobacterium-containing PbEGF1 or PbEGF1-HIGS again at 4dpi. Green fluorescence was detectable within the primary zoosporangia at 7 dpi. The white arrows indicate the primary zoosporangium of *P. brassicae.* Seedlings overexpressing PbEGF1 were designated as PbEGF1-O, whereas those inoculated with *P. brassicae* were referred to as PbEGF1-O+P. br. Similarly, seedlings expressing the PbEGF1-HIGS vector were labeled as PbEGF1-H, whereas those seedlings inoculated with *P. brassicae* were designated as PbEGF1-H+P. br.

To investigate the virulence function of PbEGF1, seedlings as previously described were inoculated with *P. brassicae*. At 10 dpi, the roots of *B. napus* were taken and subjected to peach dye staining for microscopic examination. The results demonstrated a significant reduction of *P. brassicae* colonization within the root hairs due to the silencing of *PbEGF1*. However, the seedlings with transient overexpression of *PbEGF1* exhibited an increased formation of primary zoosporangia within their root hairs, compared to the control group (Fig. 6B), the presence of some empty zoosporangia suggested the release of secondary zoospores. *Pbactin* was used for quantifying *P. brassicae* content via qRT-PCR. The results indicated that overexpression of PbEGF1 significantly enhanced the proliferation of *P. brassica*e, whereas silencing *PbEGF1* attenuated the pathogen’s accumulation (Fig. 6C). Additionally, qRT–PCR confirmed that *PbEGF1* transcript levels increased in seedlings with transient overexpression and significantly decreased in seedlings with *PbEGF1* gene silencing induced by the host (Fig. 6D).

**Fig. 6.**
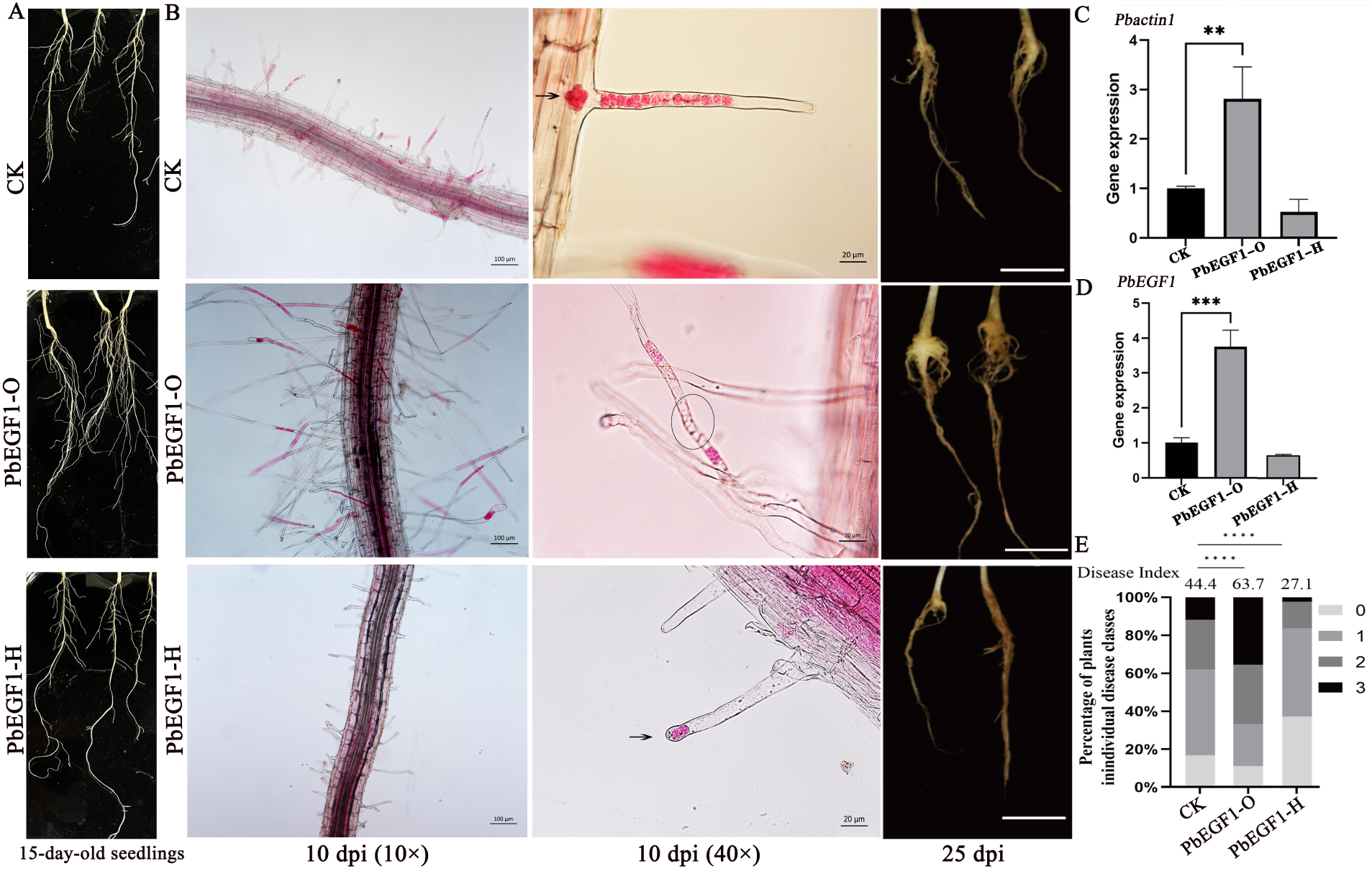
PbEGF1 enhances *P. brassicae* infection in *B. napus*. A, Root growth was measured at 15-day-old seedlings following transient overexpression of PbEGF1 and PbEGF1-HIGS. B, Representative images of root hairs and epidermal cells infected with *P. brassicae* at 10 dpi, as well as symptoms at 25 dpi, are presented. Transient overexpression of PbEGF1 containing signal peptide (PbEGF1-O) and host-induced gene silencing of PbEGF1 (PbEGF1-HIGS) were performed. C, qRT-PCR analysis was conducted to measure the transcript levels of *Pbactin1* at 10 dpi. D. qRT-PCR analysis was conducted to measure the *PbEGF1* transcript levels at 10 dpi. The control treatment was used for comparison. The experiment was repeated three times for statistical significance assessment using one-way ANOVA followed by Tukey’s multiple comparison test (**p<0.01, ***p<0.001, ****p<0.0001). E, Disease index was determined for *B. napus* plants inoculated with *P. brassicae* using a sample size of 50-60 seedlings per treatment group for Student’s t-test ( ****p< 0.0001).

The disease incidence was determined at 25 dpi. The data demonstrated that PbEGF1 facilitated *P. brassicae* infection, resulting in more severe clubroot symptoms compared to the control. However, the symptoms were alleviated upon PbEGF1 silencing (Fig. 6B). The disease index for the control group was 44.4, which increased to 63.7 in the PbEGF1 overexpression group, and was lowest at 27.1 in the host-induced PbEGF1 silenced group (Fig. 6E). These results indicate that PbEGF1 plays a crucial role in the pathogenicity of *P. brassicae*.

### PbEGF1 facilitates the proliferation of lateral roots and root hair

To elucidate the possible reason underlying PbEGF1-mediated enhancement of *P. brassicae* infection, we conducted an analysis of root growth. Overexpression of PbEGF1 in the host *B. napus* significantly enhanced the abundance of lateral roots compared to the control group (Fig. 6A). Microscopic examination revealed a substantial increase in root hair length at 10 dpi (Fig. 6A), suggesting that PbEGF1 may facilitate *P. brassicae* infection by promoting the lateral root development and elongation of root hairs, thereby expanding our understanding of its contribution to pathogenesis.

### PbEGF1 interacts with BnNHL13 protein in B. napus

Although SOBIR1 is required for cell death induction by PbEGF1 in *N. benthamiana*, no direct interaction between PbEGF1 and SOBIR1 has been detected. To elucidate the virulence mechanism of PbEGF1, we conducted yeast two-hybrid screens utilizing a *B. napus* root cDNA library infected with *P. brassicae* to identify potential interaction protein. NDR1/HIN1-like protein 13 (BnNHL13) consistently emerged across three independent screens, thus confirming its role as an interaction protein of PbEGF1.

The interaction between PbEGF1 and BnNHL13 was validated using three distinct assays. Initially, the yeast two-hybrid assay was performed with PbEGF1 as the prey protein, demonstrating a direct interaction between PbEGF1 and BnNHL13 (Fig. 7A). Subsequently, we conducted co-immunoprecipitation (Co-IP) assays using GFP-tagged PbEGF1 and FLAG-tagged BnNHL13 to validate their interaction in *N. benthamiana*. The Co-IP results demonstrated the immunoprecipitation of BnNHL13 with PbEGF1 (Fig. 7B). Finally, a luciferase complementation (Luc) assay was conducted. PbEGF1 was fused with the NLuc vector, while BnNHL13 was fused with the CLuc vector. An expression vector containing the luciferase gene served as the positive control. Transient expression was achieved through *Agrobacterium*-mediated in *N. benthamiana*. The fluorescence intensity was qualitatively assessed using a plant living-cell molecular imaging system (Fig. 7C). The results clearly demonstrated an interaction between PbEGF1 and BnNHL13.

**Fig. 7.**
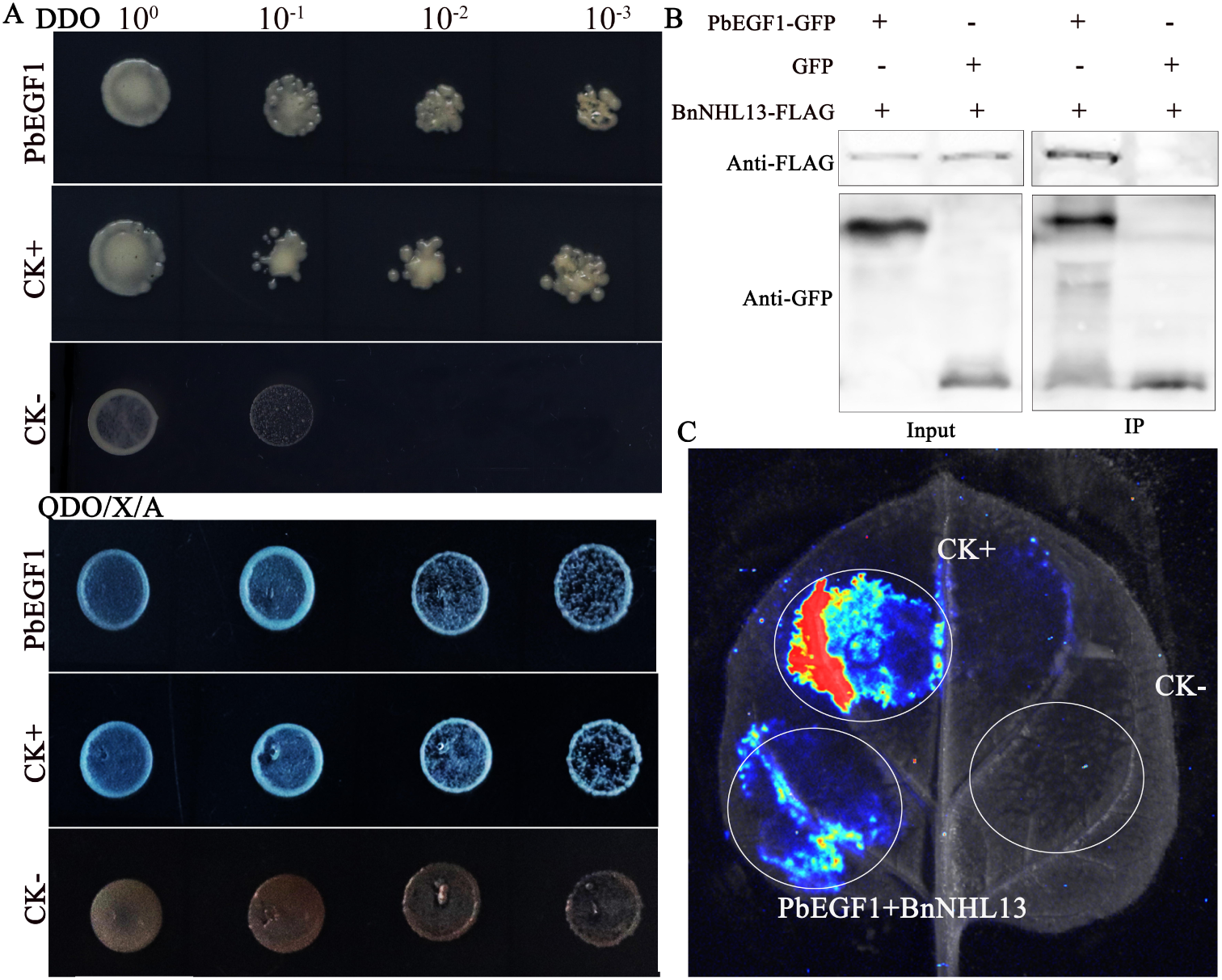
A specific interaction was identified between PbEGF1 and BnNHL13. A, Interactions between PbEGF1 and BnNHL13 were detected using a bait vector, pGADT7-BnNHL13, and a prey vector pGBKT-PbEGF1. Haploids carrying pGADT7-BnNHL13 and pGBKT-PbEGF1 were mated, and the diploids were assessed for growth in SD/-Leu/-Trp (DDO) medium as well as SD/-Ade/-His/-Leu/-Trp/X-α-Gal/AbA (QDO/X/A) medium. pGBKT7-53 and pGADT7-T was used as positive control, while the negative controls were represented by pGBKT7-Lam and pGADT7-T. B, Co-expression proteins of BnNHL13-FLAG and PbEGF1-GFP in *N. benthamiana* leaves were achieved, followed by immunoprecipitation using GFP trap beads. Western blotting analysis confirmed the co-precipitation of GFP tagged PbEGF1. C, Luciferase complementation imaging analysis was conducted on *N. benthamiana* leaves expressing PbEGF1-nLuc and BnNHL13-cLuc constructs to assess their interaction. An expression vector containing the luciferase gene was used as a positive control, while PbEGF1-nLuc and cLuc combination acted as a negative control.

### Overexpression of BnNHL13 confers enhanced resistance in plants against P. brassicae infection

To investigate the influence of the PbEGF1 target protein BnNHL13 on *P. brassicae* infection, transient overexpression and silencing of *BnNHL13* were performed in *B. napus*. The expression level of *BnNHL13* was quantified using qRT-PCR (Fig. 8D). Upon *BnNHL13* silencing, elongated root hairs and an increased number of zoosporangia were observed at 10 dpi compared to the control group (Fig. 8A). Symptoms became apparent at 25 dpi, and BnNHL13 overexpression alleviated clubroot symptoms (Fig. 8B and 8C). The expression level of *Pbactin* was induced in *BnNHL13* overexpression seedlings, whereas it was elevated in *BnNHL13*-silenced seedlings compared to the control, suggesting an increased content of *P. brassicae* in these plants (Fig. 8D). Interestingly, it was found that the overexpression of *BnNHL13* exerted an inhibitory effect on the expression of *PbEGF1*, while the silencing of *BnNHL13* enhanced the expression of *PbEGF1*, suggesting an antagonistic relationship between BnNHL13 and PbEGF1 (Fig. 8D). These findings suggest that BnNHL13 confers a certain degree of resistance against *P. brassicae* infection, whereas its silencing promotes host susceptibility to the pathogen.

**Fig. 8.**
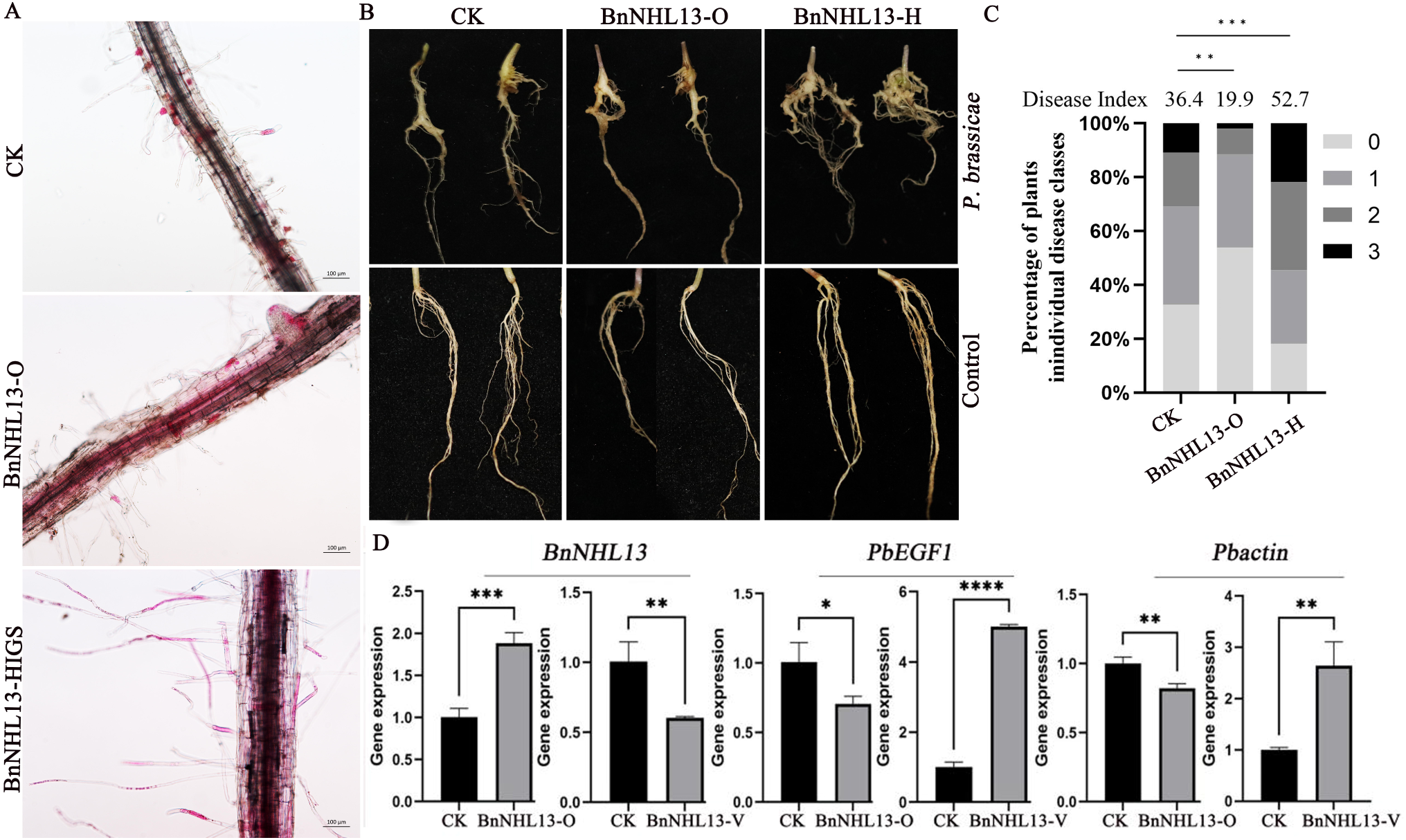
Silencing of *BnNHL13* enhances the susceptibility of *B. napus* to *P. brassicae*. A, Representative images of root hairs and epidermal cells upon infection with *P. brassicae* at 10 dpi. B, Disease symptoms are recorded at 25 dpi. C, The disease severity index of *B. napus* inoculated with *P. brassicae* at 25 dpi. The analysis was involved a minimum of 50 seedlings in each treatment group. Significant difference was performed using Student’s t-test (**p<0.01, ***p<0.001). D, qRT-PCR analysis was performed to measure the transcript levels of *Pbactin*, *PbEGF1* and *BnNHL13* at 10 dpi. *Bnactin* in *B. napus* was used as an internal reference gene. BnNHL13-O refers to transient overexpression of *BnNHL13*, BnNHL13-V represents host-induced gene silencing of *BnNHL13*. The experiment was repeated three times, and statistical significance was evaluated via one-way ANOVA followed by Tukey’s multiple comparison test (*p<0.05, **p<0.01, ***p<0.001, ****p<0.0001).

### PbEGF1 caused the degradation of BnNHL13 without affecting its subcellular localization

To investigate the underlying mechanism by which PbEGF1 interacts with BnNHL13 to effect *P. brassicae* infection, we transiently expressed BnNHL13 alone or in combination with PbEGF1 in *N. benthamiana*. Subsequently, the impact of PbEGF1 on the subcellular localization of BnNHL13 was investigated (Fig. 9A). The findings demonstrated that PbEGF1 did not alter the cellular localization of BnNHL13.

**Fig. 9.**
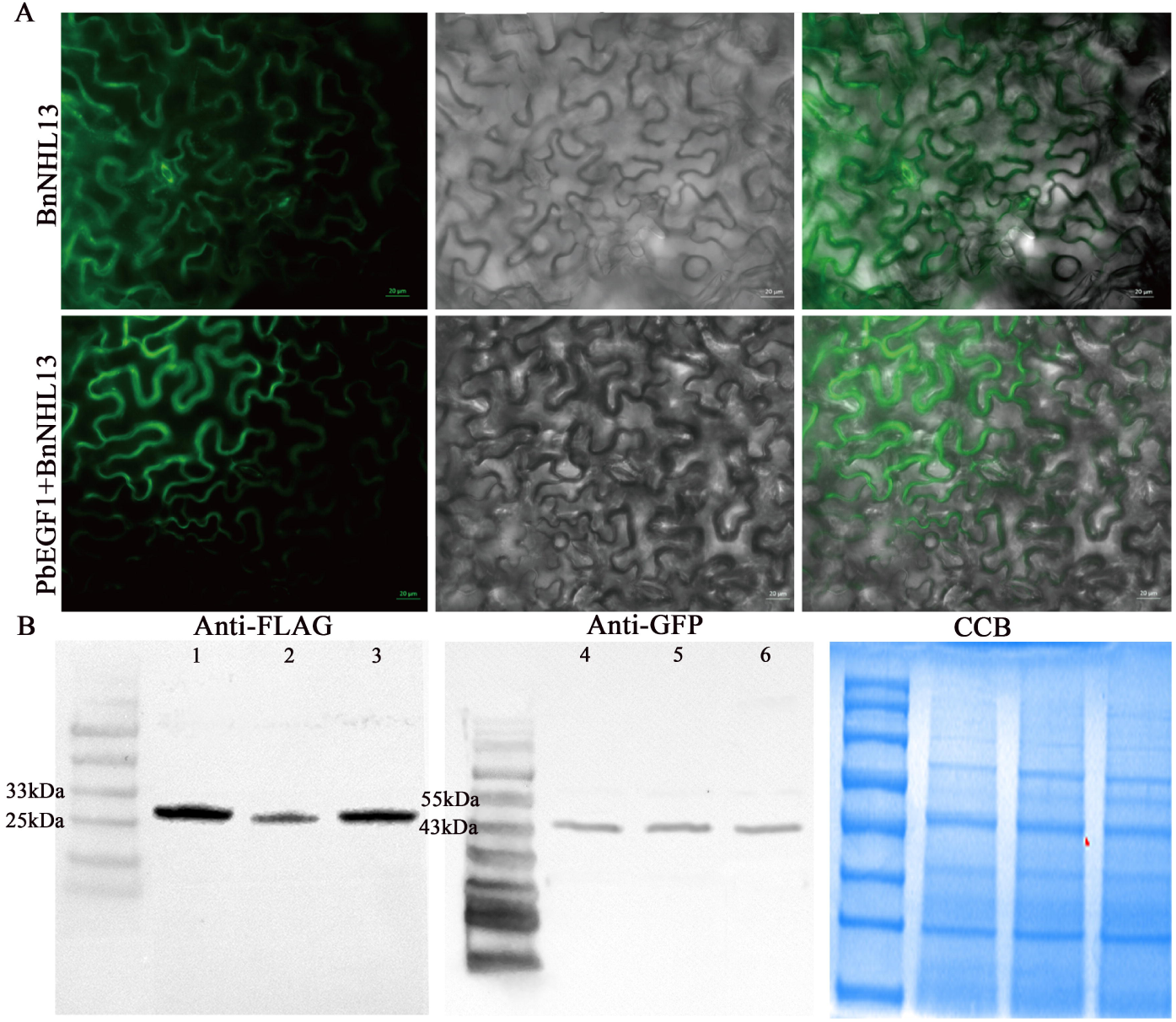
Interference effect of PbEGF1 on the host BnNHL13 was investigated. A, Transient expression of *BnNHL13* in *N. benthamiana* leaves was performed either alone or in combination with PbEGF1. The spatial distribution of the green fluorescence was examined using a fluorescence microscope at 2 days after infiltration. B, Western blotting analysis was conducted to detect the protein expression levels of PbEGF1 and BnNHL13 at 2 days post-infiltration. One day after treatment with the protease inhibitor Mg132, BnNHL13 and PbEGF1 were co-infiltrated into *N. benthamiana* leaves. Sample 1 represents the expression of BnNHL13 alone, samples 2 and 5 denote the co-expression of BnNHL13 and PbEGF1, samples 3 and 6 represent the co-expression after Mg132 treatment; sample 4 corresponds to the expression of PbEGF1 alone. BnNHL13 was fused with a FLAG tag, whereas PbEGF1 was fused with a GFP tag. Samples 1, 2 and 3 refer to FLAG antibody detection, samples 4, 5 and 6 refer to GFP antibody detection.

Furthermore, we separately infiltrated PbEGF1 and BnNHL13 into *N. benthamiana leaves*. The co-infiltration of PbEGF1 and BnNHL13 led to a significant decrease in the protein expression levels of BnNHL13 compared to those with BnNHL13 infiltration alone, indicating that PbEGF1 may facilitate the degradation of BnNHL13. Additionally, treatment with Mg132, a 26S protease inhibitor, effectively inhibited the degradation process and restored expression levels similar to those for BnNHL13 infiltration alone (Fig. 9B). These results suggest that PbEGF1 promotes the degradation of BnNHL13 through the 26S proteasome pathway.

## Discussion

Characterizing virulence effectors is essential for unraveling the pathogenic strategy. *P. brassicae* is an obligate parasitism. The manipulation of gene overexpression or knockout within its genome presents significant challenges. *Agrobacterium*-mediated transient heterologous overexpression or gene silencing in *N. benthamiana* is a widely employed method. Consequently, take into consideration the ability of *Agrobacterium-*mediate gene expression in plants, we have developed methods for instantaneously expressing or silencing *P. brassicae* genes in host rape.

Host-induced gene silencing (HIGS) is an advancement of the virus induced gene silencing (VIGS) technique. If host cells have dsRNA that is homologous to the pathogen’s sequence, it has the potential to induce gene silencing of the corresponding target (Panwar *et al*., 2018; Chen *et al*., 2022). Here, a recombinant vector containing a hairpin sequence of PbEGF1 was transiently expressed in *B. napus* via *Agrobacterium*-mediated. qRT-PCR confirmed the downregulation of PbEGF1. Fluorescence observation under microscope was also reconfirmed their successful heterologous expression in *B. napus*.

Despite the use of transient expression, the gene of *P. brassicae* could be silenced in infected hosts. Previous study employed two transient methods, host-induced gene silencing (HIGS) and spray-induced gene silencing (SIGS), to suppress the pathogenic development (Li *et al*., 2022). Otherwise, the naked or nanomaterial-bound dsRNA is prayed onto plants to induce targeted suppression of pathogenicity genes in *Magnaporthe oryzae* (Sarker *et al*., 2021). These results suggest that even though dsRNA is not stably produced, it can still silence in the host. However, the efficiency of RNA uptake by fungal pathogens is crucial for targeted gene silencing, showing variation among diverse phytopathogens and their specific genes (Sarker *et al*., 2021). The primary zoospore and zoosporangium of *P. brassicae* lack cell wall and can obtain host nutrition through phagocytosis. The green fluorescence observed in the zoosporangium indicates that the transient expression products can be uptake by *P. brassicae*, suggesting that small dsRNA molecules are also capable of being engulfed. HIGS can serve as a practical tool for investigating virulence genes in *P. brassicae*.

The heterologous expression of PbEGF1 in *B. napus* was achieved through *A. tumefaciens*-mediated, which resulted in a significant enhancement of root hair elongation and an increased quantity of *P. brassicae* infection. Recent research have elucidated that primary infection, namely root hairs infection, exhibits a critical influence, challenging the view that secondary infection is predominant (McDonald *et al*., 2014). Our findings also validate the presence of effectors in *P. brassicae* that facilitate root hair infection. The augmentation in root hair infection further exacerbates clubroot symptoms, providing supplementary evidence for the involvement of root hair infection in symptom development.

To explore the mechanism of PbEGF1 on the host, we conducted a targeted screen and successfully identified the host protein BnNHL13 as a target of PbEGF1, which is a plasma membrane-localized protein that shares similarity with NDR1/ HIN1-like proteins. NDR1 plays a broad role in host physiology and stress-induced signaling (Coppinger *et al*., 2004; Gao *et al*., 2011). *Arabidopsis* mutant *ndr1-1* displays accelerated development (Dhar *et al*., 2019). In this study, it was found that the root hair of plants with silenced BnNHL13 significantly elongated under *P. brassicae* infection. Subsequent investigations have revealed that PbEGF1 can degrade BnNHL13, suggesting that PbEGF1 promotes root hair growth and establishes conditions conducive to its own infection by possibly inhibiting BnNHL13 expression.

In order to establish a successful infection, *P. brassicae* must either fail to elicit host defense responses or suppress host resistance. NDR1 has been demonstrated to be indispensable in various plant-pathogen interaction systems (Coppinger *et al*., 2004; Gao *et al*., 2011). For instance, the absence of functional NDR1 in both Arabidopsis and cotton plants renders them more susceptible to pathogen *Verticillium*. Additionally, silencing of GmNDR1 enhances soybean susceptibility to *P. sojae* (Gao *et al*., 2011). Our findings indicate that BnNHL13 plays a role in plant resistance against *P. brassicae*. Silencing BnNHL13 enhances susceptibility to *P. brassicae,* and meanwhile stimulates root growth. Considering its multifaceted functions in plant growth and defense responses, it is not surprising that NDR1 has emerged as an important target for virulence effectors secreted by diverse pathogens. Previous reports have reported that the *Phytophthora* effector Avh241 interacts with host NDR1-like proteins, potentially disrupting the self-association of GmNDR1 and subsequently manipulating plant immunity (Yang *et al*., 2021). We observed that PbEGF1 resulted in a significant reduction in protein expression levels. Consequently, it is plausible to suggest that PbEGF1 attenuates the BnNHL13-mediated defense pathway. These findings suggest that effectors released by different pathogens can interfere with NDR1 through various ways.

The candidate effectors were transiently expressed in the model plant *N. benthamiana* for functional analysis, which is a widely employed strategy to investigate the functions of many pathogen effectors (Li *et al*., 2015b; Ma *et al*., 2015, Hu *et al*., 2019; Yang *et al*., 2021). In plant-pathogen interactions, the apoplastic space serves as a multifaceted battleground involving diverse types of interactions, determining the overall outcome of infection (Doehlemann and Hemetsberger, 2013; Ma *et al*., 2015). Here, we have identified a participant in this battlefield, PbEGF1. It elicited cell death in *A. thaliana*, *B. rapa*, and *N. benthamiana* leaves. This finding implies a potential impact on its conserved plant target. Some studies have pointed out that the induction of cell death is important for the generation of nutrient sources prior to haustoria development, or facilitating the disruption of physical barriers that promote invasion into host tissues (Ma *et al*., 2015). Previous studies have revealed that the damage of cell walls is a characteristic feature observed in susceptible hosts of *P. brassicae*, while resistant hosts do not exhibit such damage. It is likely that the effectors inducing cell death are likely to possess additional influence on host.

BAK1 and SOBIR1 are co-receptors of different PRRs, which can recognize PAMPs (Thomas *et al*., 2014; Liebrand *et al*., 2014). Certain effectors, such as XEG1 and AEP in *P. sojae*, induce cell death in *N. benthamiana* through a BAK1-dependent defense (Ma *et al*., 2015; Xu *et al*., 2021). However, PbEGF1-triggered cell death occurs independently of BAK1 but relies on the SOBIR1. SOBIR1 specifically functions in receptor complexes containing LRR-RLPs. PbEGF1 detection was mediated by a cell surface pattern recognition receptor. Otherwise, *P. brassicae* effector Pb257 also elicits cell death in *N. benthamiana* by depending on SOBIR1 (Yang *et al*., 2024). We propose that PbEGF1 and Pb257 may target distinct receptor-like proteins (RLPs) to modulate the same defense signal pathway. These findings indicate that pathogen effectors target diverse host factors to disrupt host defense. Pathogens secrete effectors to manipulate or evade host defense responses, thereby facilitating successful establishment of infection.

### Conclusions

In this study, the *P. brassicae* effector PbEGF1 and its target protein BnNHL13 in *B. napus* were identified. PbEGF1 can interfere with BnNHL13, promoting root hair elongation and primary infection, thereby enhancing host susceptibility. This study firstly pointed out the role of the effector in the primary infection of *P. brassicae* and further confirmed the significance of root hair infection in symptom formation. Additionally, a host-induced gene silencing method was established using agrobacterium-mediated, which not only provided a method to investigate gene function in *P. brassicae* but also offered a new strategy for preventing clubroot.

## Supplementary data

The Supplementary table 1. The primers sequence.

## Acknowledgments

We would like to give great thanks to Dr Wenming Wang and Dr Jing Fan from Sichuan Agriculture University for kindly providing the vectors.

## Author contributions

HY and JBD: conceptualization, funding acquisition and writing draft. YSX and YSZ: experiment conduction. HY, YSX and YPS: formal analysis. All authors have read and approved the manuscript.

